# Immune stromal components impede biological effectiveness of carbon ion therapy in a preclinical model of pancreatic ductal adenocarcinoma

**DOI:** 10.1101/2025.05.17.654632

**Authors:** Katy L. Swancutt, Eslam A. Elghonaimy, James H. Nicholson, Laurentiu M. Pop, Brock J. Sishc, Elizabeth M. Alves, Cassandra Hamilton, Adam Rusek, Anthony J. Davis, Raquibul Hannan, Michael D. Story, Todd A. Aguilera

## Abstract

The tumor landscape of pancreatic ductal adenocarcinoma (PDAC) is refractory to conventional photon radiotherapy (RT) due to a fibrotic tumor microenvironment (TME) that promotes chronic hypoxia and reduced immune surveillance. The radiobiological factors unique to carbon ion radiotherapy (CIRT), such as high linear energy transfer (LET) and less dependence on oxygen, make it well-suited to overcome the PDAC TME. Here, we utilized clonal syngeneic KPC pancreatic tumor cell lines and tumors to examine this postulate and to identify underlying factors that impact the response of PDAC to CIRT. While KPC cell lines exhibited radiobiologic effectiveness (RBE) greater than 3, subcutaneous tumors in the mouse hind leg showed lower RBEs – 1.3 based on quintupling time – at a LET of 75 keV/μm. Four days after CIRT, we observed widespread transcriptomic changes in the tumor immune microenvironment (TME), suggesting increased infiltration of anti-tumor immune cells, elevated expression of anti-tumor T cell cytokines, MHC class I molecules, and co-stimulatory signals. Fewer immunologic changes were observed following photon irradiation. By seven days after CIRT, tumor-supportive transcriptomic programs characterized by pro-tumor cytokines, M2 macrophages, and cancer-associated fibroblasts (CAFs) emerged, promoting resistance and limiting the durability of tumor growth delay. These findings suggest that CIRT may offer a favorable platform compared to conventional photon radiation therapy for combining with immunotherapies. Furthermore, these data highlight the risk of using *in vitro* survival data alone in treatment planning and indicate that underlying TME factors impact the response of PDAC *in vivo*.

## Introduction

Treatment of pancreatic ductal adenocarcinoma (PDAC) is complicated by a lack of early-stage detection, a propensity toward local invasiveness, proximity to adjacent organs at risk, and the ability to metastasize [1]. The use of conventional photon radiation therapy (RT) to treat PDAC in the neoadjuvant setting or for consolidative treatment of unresectable disease is associated with improved local control; however, in a large-scale, randomized, Phase III clinical trial chemoradiotherapy (CRT) failed to improve overall survival compared to chemotherapy alone [2]. This suggests that conventional photon RT is not expected to improve survival in the absence of treating a low-risk population or excellent systemic therapies given the competing risk of metastatic progression. However, improved local control could improve survival with effective systemic immune surveillance or optimal patient selection.

Carbon ion radiation therapy (CIRT) offers distinct advantages for treating pancreatic ductal adenocarcinoma (PDAC) due to its unique physical and biological properties. Unlike photons, carbon ions deliver most of their energy within the spread-out Bragg Peak (SOBP), enabling highly conformal dose delivery with minimal entry and no exit dose—sparing nearby organs like the stomach and duodenum. Within the SOBP, carbon ions deposit energy with high linear energy transfer (LET), causing dense, complex DNA damage that requires multiple DNA repair pathways to resolve, leading to enhanced cell killing [3–7]. Further, the increased damage resulting from direct ionization events reduces the dependence of damage fixation via oxygen, making CIRT well-suited for hypoxic, desmoplastic tumors like PDAC [8–10]. Due to these physical and biological properties, carbon and other high-LET ions have a higher relative biological effectiveness (RBE), requiring lower absorbed doses for the same biological effect as photons [8, 10, 11]. However, translating CIRT clinically requires accurate RBE modeling, which is complicated by variability in biological systems, dosimetry, and LET distribution [12].

Importantly, CIRT may also elicit distinct immunogenic effects, enhancing its potential when combined with immunotherapy [13].

In clinical trials comparing chemotherapy vs photon CRT, median survival was found to be 15.2 vs 16.5 months, respectively [2], while dose-escalated RT achieved 18.4 to 24+ months [14, 15]. By way of comparison, early phase clinical trials in Japan found CIRT to be safe and effective in resectable PDAC [16] and when combined with chemotherapy, in unresectable PDAC [17–19].The median survival of CIRT with chemotherapy was 19.6 months in early trials and 21.5-25.1 months in later studies [18, 19]. CIRT for recurrent PDAC was also promising [19, 20]. However, progress beyond Phase I/II trials has been hindered by administrative challenges (NCT01795274) and poor patient accrual (NCT03822936) [21, 22]. However, results from other Phase I carbon clinical trials, recruiting (NCT04082455) or complete (NCT03949933 and NCT03403049), are pending. Given the promising results seen from Japan that suggest that CIRT may provide unique advantages over conventional photon or chemoradiotherapy suggests that understanding the pancreatic tumor microenvironment may help determine whether CIRT truly offers unique advantages over photon RT.

In this study, we investigated the differential effects of photon RT and CIRT in syngeneic, poorly immunogenic KPC (Cre driven Kras and TP53) pancreatic cancer mouse models. After establishing parameters to evaluate immune responses to photon RT and immunotherapy, we observed a discrepancy between *in vitro* and *in vivo* RBE. We hypothesized that TME components reduce photon RT effectiveness and contribute to CIRT resistance. We performed comprehensive bulk RNA sequencing to identify radiation-induced response pathways. CIRT resulted in a robust immune activation by day four, with increased immune infiltrates, cytokine signaling, cytotoxic T cells, MHC class I, and co-activation markers, exceeding changes seen with photon RT. By day seven, there was an immunosuppressive anti-tumor response after CIRT, marked by pro-tumor cytokine expression and expansion of M2 macrophages and CAFs. These findings reveal TME-driven limitations to CIRT efficacy and support the development of rational and temporal CIRT-immunotherapy combinations and may highlight the importance of the immune response in clinical RBE determinations as opposed to the use of *in vitro* RBE values as predominant influencers in treatment planning.

## Materials and Methods

### Animals

KPC mice, backcrossed to C57BL/6, were bred from founders gifted by David DeNardo (Washington University). They carried LSL-KrasG12D, heterozygous p53 loss, and p48 (*Ptf1a*)-driven cre recombinase. Female 6–8-week-old C57BL/6 mice were purchased from Charles River and Jackson Laboratories. Animal studies were approved by UT Southwestern and, for carbon ion and control experiments, by BNL IACUC. Groups consisted of five mice for abscopal and transport studies, and nine to ten mice for carbon ion and photon experiments.

### Tumor Model

Clonal cell lines were derived from tumors of KPC mice, selected over KPPC mice (homozygous p53^flox/flox^) for slower tumor progression (median survival: 28.5 vs. 10 weeks; Supplementary Figure 1A). Tumors were minced and digested for 40 min at 37°C with 20 μg/ml liberase TL (Sigma #5401020001) and 400 μg/ml DNAse I (Sigma #D5025-150KU), then quenched with DMEM (Fisher #MT10013CV) containing 10% HyClone FetalClone II Serum (Cytiva #SH30066.03) and filtered through a 70 µm strainer. Cells were cultured until adipocytes and fibroblasts were no longer detected microscopically. Single-cell clones were expanded after plating 2 cells/well, verifying only one visible colony per well, that resulted in over 60 clonal lines. Clones were confirmed negative for mycoplasma by PCR (ATCC Universal Mycoplasma Detection Kit #30-1012K). For abscopal experiments, 1×10⁶ KPC36 cells in 1:1 PBS:Matrigel (Corning #356234) were injected bilaterally; for KPC63, 1×10⁶ cells were injected on the right and 5×10⁵ on the left. For single-tumor studies, 8×10⁵ KPC36 cells in 1:1 PBS:Matrigel were injected into the right hind leg. Tumors were measured with calipers and volume calculated as longest length x longest length x width/2. Mice were stratified into groups once tumors reached 100 ± 15 mm². Carbon ion beam time was divided based on availability across four non-consecutive days. In a pilot experiment, we verified that tumor growth was unaffected by shipping, consistent with findings from Brownstein *et al.* (Supplemental Figure 3B)[23], and later confirmed that travel and anesthesia had no impact on tumor growth using untreated controls (Supplemental Figure 3C). Mice were housed at Brookhaven for over three days to acclimate before initiating treatment.

### Antibodies

Therapeutic and isotype control antibodies were purchased from BioXCell and administered intraperitoneally in PBS (200 μL total volume). Anti-mouse αCTLA-4 (clone 9D9, #BE0164), αPD-1 (clone RMP1-14, #BE0146), and their isotype controls (mouse IgG2b, #BE0086; rat IgG2a, #BE0089) were given at 200 μg per dose. αCD40 (clone FGK4.5/FGK45, #BE0016-2) and its isotype control (rat IgG2a, #BE0089) were administered at 100 μg per dose.

### Single Cell RNA Sequencing

KPC63 tumors developed from 1×10⁶ cells in 1:1 PBS:Matrigel injected subcutaneously into the hind leg. Tumor-draining lymph nodes (TDLN) were collected from three mice per treatment group. To obtain single-cell suspensions, TDLNs were homogenized gently and filtered through a 40 μm filter in 10% serum-supplemented medium. Tumor tissue was digested in 5 mL HBSS (Fisher #MT20023CV) and RPMI with L-glutamine (Fisher #MT10041CM) (4:1), supplemented with 3% FetalClone II, 10 μg/ml DNAse I, 10 μM Rho/ROCK inhibitor Y-27632 (Sigma #Y0503-5MG), and 100 U/ml Collagenase IV (Worthington #LS004188) for 45 min at 37°C with shaking at 50 RPM. After digestion, serum-supplemented medium was added, and cells were filtered through a 40 μm filter. The cell suspension was pelleted, RBCs lysed in ACK buffer for 5 min at room temperature, and cells resuspended in FetalClone II containing 10% DMSO (Fisher #BP231-1) before freezing at −80°C overnight and storing in liquid nitrogen.

Cells were slowly revived by addition of 1 mL RPMI with 5% FetalClone II media per minute until thawed. After counting using a TC20 automated cell counter (BioRad), cells were adjusted to 1000 cells/μL in 0.04% BSA/PBS (Fisher #BP9706100, #MT21040CV), and 1 mL/sample was submitted to the Next Generation Sequencing Core, UTSW. Cells were loaded onto a 10x Chromium microfluidic chip using the single cell 3′ kit (V3 chemistry) to capture 10,000 cells. Library construction and all subsequent procedures followed the manufacturer’s standard protocol. Single-cell libraries were sequenced via NovaSeq (Illumina) to a depth of 50,000 reads per cell.

### Single Cell RNA Sequencing Data Processing

Raw data and FASTQs were generated by the Next Generation Sequencing Core, UTSW. Using Cell Ranger 5.0.1, FASTQ files were aligned to the mouse genome (mm10) with STAR 2.7.2a and counted using default parameters. The matrix was imported into Seurat (3.2.3) via R (4.0.3) for quality control and analysis. Cells with <200 genes, genes in <3 cells, or >20% mitochondrial genes were excluded. Data were normalized with NormalizeData, and variable genes detected with FindVariableFeatures. Cell cycle scores were calculated by CellCycleScoring, with the S phase score subtracted from the G2M score. Data were scaled using linear regression on counts and cell cycle differences. PCA was performed with RunPCA, and batch effects corrected with Harmony via SeuratWrappers. Clusters were identified using FindNeighbors, FindClusters (resolution 0.5), and UMAP. A FindAllMarkers table was created, and clusters were defined using SingleR, celldex with ImmGenData (https://www.immgen.org/), and canonical markers (Supplementary Figure 2B).

Cell population enrichment across treatment groups was assessed using the Ro/e ratio (observed/expected cell number) as previously described [24, 25], with chi-squared tests (chisq.test) to identify enriched cell subtypes (Ro/e >1). For TME classification, cellular modules were adapted from George *et al*. (2024), converting to mouse genes and calculating scores with AddModuleScore in Seurat. Full lists of genes used to define response and resistance programs are provided in Supplementary Table 1. Radiation response was assessed via DNA damage repair genes, hypoxia markers, pro-inflammatory genes, and TME remodeling factors. Radiation resistance was analyzed using EMT signature genes, ROS scavenging factors, and fibrotic TME markers [26]. Immunotherapy response was evaluated based on cytotoxic T cell response, type I IFN response, and DC activation markers, while resistance was characterized by T cell exhaustion genes, MDSC immunosuppression markers, and fibrotic TME factors. Targetable genes were adapted from George *et al*. (2024) and https://clinicaltrials.gov/.

### Immunoblotting

One million cells per clone were lysed on ice in RIPA buffer (EMD Millipore #20-188) supplemented with protease inhibitors, phosphatase inhibitors, DTT, and PMSF (Sigma #PPC1010, #10708984001, and #10837091001). Protein concentration was measured using Bradford reagent (BioRad #5000205). Protein lysates were mixed with 2X Laemmli sample buffer (BioRad #1610737) and boiled at 100°C for 10 minutes. 30 μg of protein per sample was separated by 10% acrylamide gel electrophoresis at 60 volts and transferred onto PVDF membranes (BioRad #1620174) at 4°C overnight at 22 volts. Transfer efficiency was confirmed by Ponceau S staining (VWR #97062-458). Membranes were blocked in 5% non-fat dry milk in TBST (Cell Signaling #9997), washed three times, and incubated overnight at 4°C with primary antibodies (1:1,000 dilution in 5% BSA/TBST): rabbit anti-STING (D1V5L #50494), rabbit anti-cGAS (D3O8O #31659), and rabbit anti-vinculin (E1E9V #13901) (all from Cell Signaling). After four washes with TBST, membranes were incubated with HRP-conjugated goat anti-rabbit secondary antibody (Jackson ImmunoResearch #111-035-003; 1:3,000 dilution in 1% milk/TBST) for 2 hours at room temperature with shaking. Detection was performed using Pierce™ ECL substrate (Thermo Fisher #32209) and developed on X-ray films (Lightlabs #X-3001) after 5 and 45 minutes of exposure.

### In Vitro Evaluation of RBE

The RBE of the KPC36 and KPC63 cell lines was determined via clonogenic survival assay [27]. The carbon ion beam (test radiation) at CNAO (Pavia, Italy) had a linear energy transfer (LET) between 74.1 and 89.3 keV/μm. Gamma radiation from a Cesium-137 irradiator (JL Shepherd and Associates, San Fernando, CA) at UTSW served as the low LET reference radiation, as previously reported [26, 27]. Survival curve fitting was performed using MATLAB with either Repairable Conditionally Repairable (RCR) modeling [28] or linear quadratic (LQ) modeling [29].

### In Vivo Photon Radiation Delivery

Tumor irradiation with photons was performed at UTSW using inhalant anesthesia (isoflurane) to immobilize mice for X-ray treatment. A 250 keV X-ray beam was delivered via an XRAD 320 device (Precision X-Ray, Inc.) with a 10 mm diameter, circular, copper collimated beam at 5 cm source to surface distance (SSD) [30]. The XRAD 320 delivers dose according to an input in units of whole seconds, making 3.36 Gy the nearest mechanically achievable dose equivalent to 3.30 Gy. Tumors were centered in the beam using a machined metal insert suspended from the collimator.

### In Vivo Carbon Ion Radiation Delivery and RBE Calculations

Tumor irradiation with carbon ions was performed at the NASA Space Radiation Laboratory (NSRL) at Brookhaven National Laboratory (BNL), Upton, NY. Mice were anesthetized with 80 mg/kg ketamine and 10 mg/kg xylazine, then positioned on a plexiglass frame with magnetic attachments to align the tumor within the collimator. Tumors were aligned with the beam using laser sights to position the tumor within a SOBP generated via a 3D-printed, acrylonitrile butadiene styrene (ABS), rotating compensator creating dose steps equivalent to 0.5 mm of HDPE. Anesthetized mice were immobilized on a custom stage to align the SQ right hind leg tumor with the collimated beam, positioning each ∼100 mm³ tumor within a 6 mm region starting 9 mm beyond the proximal edge of the SOBP. A total dose of 3.30 Gy at an average LET of 75 keV/μm was delivered in one fraction. Mice were reversed from anesthesia as needed with atipamezole. Necropsies for tissue collection were done at BNL, while a subset of irradiated and control mice was shipped to UTSW for further tumor growth measurement.

These mice were quarantined, tested for common murine pathogens, and found to be pathogen-free.

Exponential growth rate used to determine tumor tripling and quintupling time was calculated using y = y_0_e^kx^, where y_0_ was the y intercept and k was the rate constant. Photon equivalent doses to achieve isoeffective tripling and quintupling times were calculated based on linear regression of time versus dose.

### RNA Extraction and Sequencing

Tumor tissue was preserved in RNAlater (Qiagen) and stored at −80°C. RNA was extracted using Qiagen RNeasy Plus Kits and quantified with Nanodrop 2000 and Agilent RNA 6000 Pico Kit on the Agilent 2100 Bioanalyzer. Samples with RNA integrity numbers (RIN) > 7.0 were deemed acceptable. RNA was prepared for sequencing using the Illumina Stranded mRNA Prep kit, starting with 25-1000 ng of total RNA for poly(A) mRNA enrichment. Enriched mRNA underwent fragmentation, cDNA synthesis, end repair, and adapter ligation, followed by PCR amplification and purification with AMPure XP beads (Beckman Coulter). Libraries were evaluated for quality using the Agilent 2100 Bioanalyzer and quantified with the Qubit dsDNA HS Assay Kit (Thermo Fisher Scientific). Sequencing was performed on an Illumina NovaSeq 6000 platform (North Texas Genome Center) with a target of 50M reads per sample. Raw reads were processed with FastQC for quality control, trimmed using Trimmomatic, and gene expression was quantified with RSEM. TPM values were imported into R via tximport and analyzed with DESeq2 for differential gene expression. Gene set enrichment analysis (GSEA) was performed using clusterProfiler with MSigDB. TME classification, and radiation and immunotherapy resistance and response were analyzed as described in the Single Cell RNA Sequencing Data Processing section, with module scores calculated using GSVA [31].

### Statistics

Tumor volume for each group of CIRT-treated animals and related photon RT or untreated controls was modeled using a repeated measures mixed model that allows for errors to be correlated within mice from SAS9.4 (SAS Institute Inc, NC). Figures were generated using GraphPad6 (Software Inc, CA).

## Results

### Characterization of carbon and photon response to pancreas cancer clones

To investigate radiation responses, we generated clonal cell lines from murine exocrine pancreatic tumors by crossing LSL-*Kras*G12D, *TP53*flox, and *p48-Cre* mice to produce triple transgenic offspring. Clones with LSL-*Kras*G12D/wt, *TP53*flox/wt, and *p48-Cre* genotypes (KPC) were established through enzymatic digestion and single-cell clonal selection (Figure 1A). To assess relative radiosensitivity, clonogenic survival assays were performed on randomly selected KPC clones (Figure 1B). After subcutaneous (SQ) transplantation in mice radiation responses varied, with some clones exhibiting ulceration prior to treatment (KPC12) or after irradiation (KPC28), while others occasionally formed large cysts (KPC62) (Supplemental Figure 1B).

**Figure 1:**
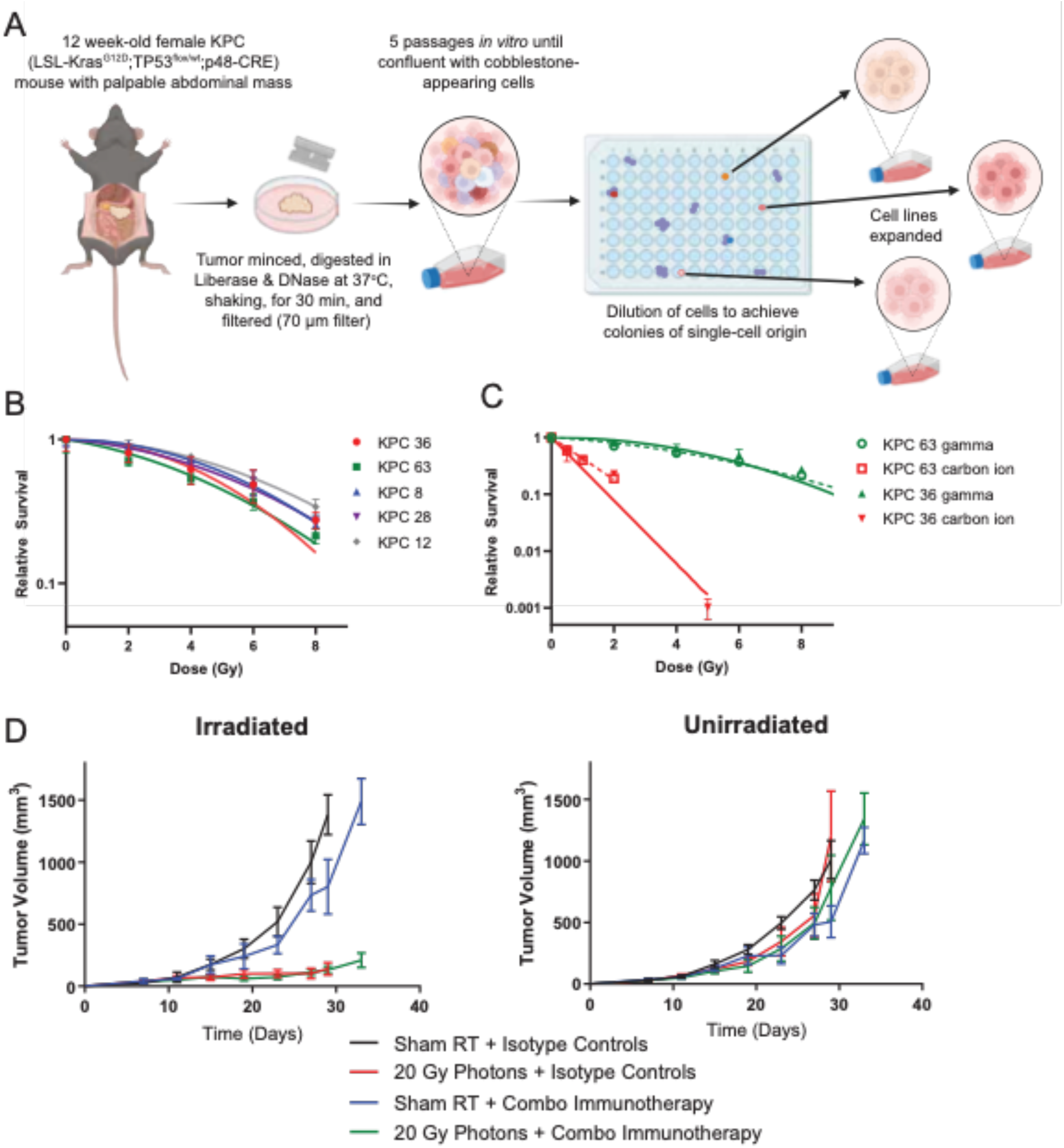
KPC tumors are resistant to photon RT and immunotherapy but carbon ions demonstrate high RBE *in vitro*. A, Expansion of multiple KPC clonal cell lines from singlecell origin. **B,** Clonogenic survival of selected KPC clones following Cesium-137 gamma irradiation. **C,** Clonogenic survival in two KPC clones treated with a carbon ions or gamma radiation. **D,** KPC36 tumors were injected subcutaneously into bilateral flanks of C57BL/6 mice (n = 5). Photon RT groups received 20 Gy to the right tumor on Day 8; combination immunotherapy groups were treated with αCTLA-4 (Days 1, 4, 7), αPD-1 (Days 8, 11, 14), and αCD40 (Day 8).

To determine the radiation sensitivity of the pancreatic cancer cells to higher linear energy transfer (LET) carbon particles, we used a clinical carbon ion beam to evaluate two clonal cell lines as previously reported, using KPC36 and KPC63, evaluated by clonogenic survival assays [27]. Our results showed that both cells exhibited enhanced sensitivity to carbon ions, characterized by the loss of the semi-log shoulder curve (Figure 1C). Furthermore, we calculated the RBE of carbon ion radiation using Reparable Conditional Repairable (RCR) modeling at an endpoint of 10% survival, yielding RBE values of 4.78 for KPC36 and 3.33 for KPC63, compared to 5.01 and 3.62, respectively, obtained from Linear Quadratic (LQ) modeling (Table 1).

**Table 1:** Surviving fraction at 2 or 3.5 Gy and RBE at surviving fraction 10% or mean inactivation dose (MID) according to either the Repairable Conditional Repairable (RCR) or Linear Quadratic (LQ) models.

| Cell Line | Model | Gamma<br>SF<sub>2 Gy</sub> | Gamma<br>SF<sub>3.5 Gy</sub> | Carbon<br>SF<sub>2 Gy</sub> | Carbon<br>SF<sub>3.5 Gy</sub> | RBE<sub>SF10%</sub> | RBE<sub>MID</sub> |
| --- | --- | --- | --- | --- | --- | --- | --- |
| KPC36 | RCR | 0.8445 | 0.6007 | 0.0777 | 0.0114 | 4.78 | 6.04 |
| KPC63 | RCR | 0.8159 | 0.5899 | 0.1873 | 0.0533 | 3.33 | 4.04 |
| KPC8 | RCR | 0.9043 | 0.7558 | --- | --- | --- | --- |
| KPC12 | RCR | 0.8979 | 0.7787 | --- | --- | --- | --- |
| KPC28 | RCR | 0.8374 | 0.6814 | --- | --- | --- | --- |
| KPC36 | LQ | 0.8930 | 0.7070 | 0.0772 | 0.0113 | 5.01 | 6.75 |
| KPC63 | LQ | 0.7984 | 0.6200 | 0.1803 | 0.0499 | 3.62 | 4.33 |
| KPC8 | LQ | 0.9202 | 0.7751 | --- | --- | --- | --- |
| KPC12 | LQ | 0.9216 | 0.7988 | --- | --- | --- | --- |
| KPC28 | LQ | 0.8721 | 0.7237 | --- | --- | --- | --- |

To explore the immunologic response to radiation, we evaluated the effects of 20 Gy photon irradiation on KPC36 and KPC63 tumors, both with and without combination immunotherapy. The immunotherapy regimen, comprising a CD40 agonist, CTLA-4, and PD-1 checkpoint blocking antibodies (Figure 1D, Supplemental Figure 1C-D), was applied to assess its impact on tumor growth on the irradiated and a contralateral unirradiated tumor [32]. Our results revealed no significant benefit with the combination immunotherapy alone, and no evidence of clinically meaningful enhanced responses with photon RT. Observed therapeutic resistance suggests that the addition of immunotherapy to photon RT may not offer significant benefits in this context.

### KPC tumors exhibit complex resistance to photon RT and immunotherapy

To elucidate the immunological mechanisms underlying treatment resistance in KPC tumors, we conducted single-cell RNA sequencing (scRNAseq) analysis. We injected KPC63 cells SQ in the hind leg of naïve mice and compared three treatment groups: 10 Gy photon RT, combination immunotherapy, or isotype control. TDLNs were harvested, dissociated into single-cell suspensions, and analyzed on day 16 post-irradiation (Figure 2A). Following integration and batch effect removal, we identified and annotated 12 distinct cellular subpopulations visualized on the t-distributed stochastic neighbor embedding (tSNE) plot (Figure 2B, Supplemental Figure 2A-B). Our analysis revealed treatment-specific alterations in immune cell composition: photon RT substantially decreased B cell populations and reduced T cell frequencies, while increasing myeloid cell populations including granulocytes, macrophages, and stromal components. In contrast, combination immunotherapy maintained T cell populations but was associated with elevated regulatory T cells (Tregs) compared to other groups (Figure 2C).

**Figure 2:**
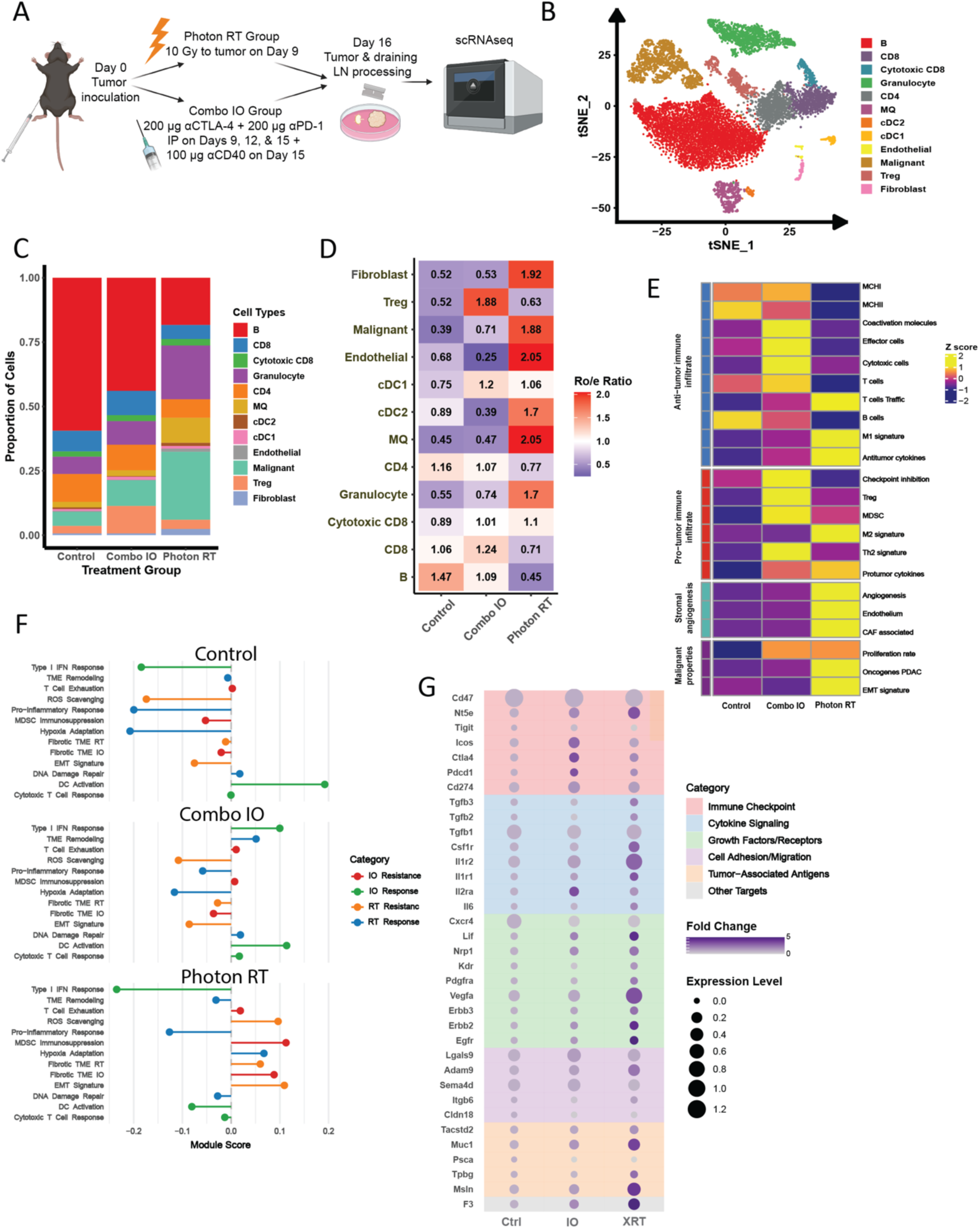
Photon RT or immunotherapy induce complex resistance mechanisms in KPC tumors. A, scRNA sequencing tissue collection and processing schema **B,** t-SNE plot of 12,661 cells in merged data. **C,** Proportion of cell types in control, and treatment groups. **D,** Heatmap of cell type enrichment scores indicated by Ro/e score. **E,** TME classification changes heatmap displays enrichment scores for gene signatures categorized into four major functional groups. D = “Depleted”, IE = “Immune enrich”, IE/F = “Immune enrich/Fibrotic”. **F,** Lollipop plot of module scores for gene signatures associated with photon RT and immunotherapy response or modules. **G,** Dot plot of targetable gene expression categorized by function.

Quantitative assessment using the Ro/e Ratio revealed distinct cellular enrichment patterns between treatments (Figure 2D). While showing a small enrichment of cytotoxic CD8 over control group, the photon RT group also showed significant enrichment of macrophages, endothelial cells, fibroblasts, and malignant cells, suggesting an immunosuppressive microenvironment. In contrast, tumors treated with combination immunotherapy exhibited a higher Treg ratio, indicative of an environment potentially more favorable to the tumor. Notably, dendritic cell (DC) populations showed a different trend: cDC2s increased after RT, whereas CD86+ DCs were enriched in the immunotherapy group, suggesting distinct effects of these treatments on immune function and tumor control.

Applying the TME classification framework [33], we observed that untreated KPC tumors and TDLN exhibited an Immune Depleted phenotype. IO treatment shifted the profile toward an Immune Enriched subtype characterized by elevated expression of both anti-and pro-tumor immune signatures without increased fibrosis or stromal components. In contrast, photon RT induced an Immune Enriched/Fibrotic phenotype, featuring upregulated expression of both anti-tumor responses and pro-tumor activities, coupled with enhanced stromal angiogenesis and malignancy-associated gene signatures (Figure 2E).

To elucidate the mechanisms underlying treatment resistance in KPC tumors, we investigated the immunologic responses to photon RT and IO. Our analysis revealed distinct patterns of gene expression changes across both treatments, highlighting the complex interplay between tumor, immune, and stromal compartments. Photon RT suppressed Type I Interferon response genes but enriched fibrosis-related pathways. In contrast, immunotherapy enriched Type I interferon, DC activation, and cytotoxic T cell response genes that were suppressed by photon RT. Notably, both treatment types induced T cell exhaustion and MDSC-mediated immunosuppression, yet the former was associated with a specific enrichment of genes involved in fibrosis unique to the photon RT group (Figure 2F, Supplemental Table 1).

Furthermore, our analysis identified distinct resistance mechanisms underlying the effects of photon RT, including suppression of TME remodeling, pro-inflammatory response, and DNA damage repair (Figure 2F). These findings underscore the diverse resistance profiles generated by different therapeutic strategies, with unique mechanistic contributions from immune, stromal, and tumor compartments.

Given the multiple mechanisms by which KPC tumors resist therapy-induced changes in the TME, we sought to identify clinically relevant targetable genes that could mitigate therapy resistance. We analyzed expression patterns of genes currently under investigation in PDAC clinical trials. This approach revealed several promising candidates for combination therapy with RT, including CD47 (expressed across multiple cell types), Vegfa (predominantly in malignant cells and granulocytes), and Il1r2 (expressed by granulocytes and cDC2 cells) (Figure 2G, Supplemental Figure 2E-F). While transcript-level changes require protein-level validation, these findings establish a foundation for evaluating CIRT’s potential to overcome the complex resistance mechanisms observed with conventional photon RT.

### Carbon ion irradiation of pancreas tumors yielded a lower-than-expected relative biological effectiveness

CIRT, characterized by high LET, is believed to be more effective in cancer therapy and may induce immune effects distinct from the mixed pro-and anti-tumor responses observed after photon RT [13]. To evaluate the potential of CIRT for tumor control in PDAC, we treated tumor-bearing mice at Brookhaven National Laboratory’s NASA Space Radiation Laboratory. A 100 MeV/n carbon ion beam and spinning compensator wheel were used, producing a Bragg peak positioned at a depth of 2.55 cm in high-density polyethylene (HDPE), followed by a steep dose drop-off (Figure 3A-D). For these experiments, KPC36 cells were chosen for injection due to their low incidence of ulceration and intact expression of cGAS and STING (Supplemental Figure 3A).

**Figure 3:**
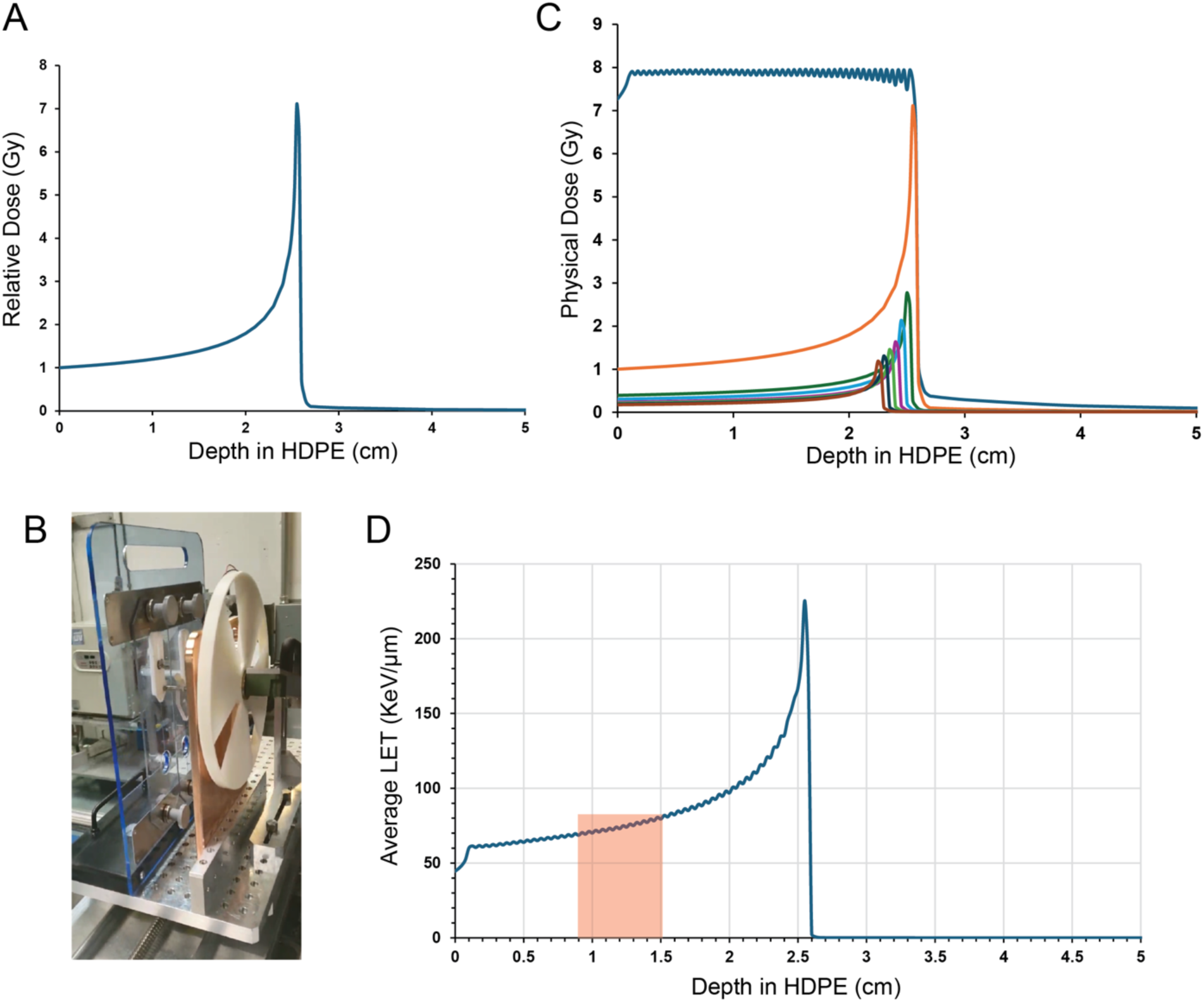
Carbon ion particle beam characteristics facilitated targeted delivery of dose to the tumor. A, Depth dose curve of the 100 MeV/n carbon ion beam with energy deposition represented as relative dose normalized to the entrance dose in HDPE (density of 0.97 g/cm3). **B,** Spinning ABS compensator wheel (density of 1.2 g/cm3) used to generate a SOBP. **C,** Monte Carlo modeling of the expected physical dose distribution, normalized to entrance dose of distal the contribution, of the SOBP using 0.5 mm step size. **D,** Simulated dose-weighted LET using 9 mm HDPE bolus before the tumor, positioned within the red highlighted region.

This modest dose was selected based on clonogenic assay results (Figure 1C) to achieve an isoeffect equivalent to 10 Gy photons. However, compared to untreated controls, 3.30 Gy carbon ions produced a less-than-expected tumor growth delay (Figure 4A). We compared treatment to 3.36 Gy photons using an XRAD 320 small animal irradiator, revealing that 3.30 Gy carbon ions induced a prolonged tumor growth delay than 3.36 Gy photons but less than that of 5 Gy photons (Figure 4B). Modeled average tumor growth curves for photon doses up to 20 Gy (Supplemental Figure 3D-E) facilitated comparison of carbon ion and photon isoeffects, including tumor growth delay and growth rate (Figure 4C-D). Exponential growth rates were similar across radiation types (Figure 4C), and tumor tripling and quintupling times for 3.30 Gy carbon ions were 12.3 ± 1.3 days and 15.8 ± 1.3 days, respectively (Figure 4D). Linear regression fitting of tripling and quintupling times versus photon dose yielded isoeffective doses of 5.49 Gy and 4.15 Gy photons, respectively (Figure 4D). The relative biological effectiveness (RBE) calculated based on tumor tripling or quintupling times was 1.7 and 1.3, respectively. Overall, the *in vivo* RBE was approximately half of that determined *in vitro*, highlighting the impact of factors such as the tumor microenvironment that may attenuate radiation sensitivity.

**Figure 4:**
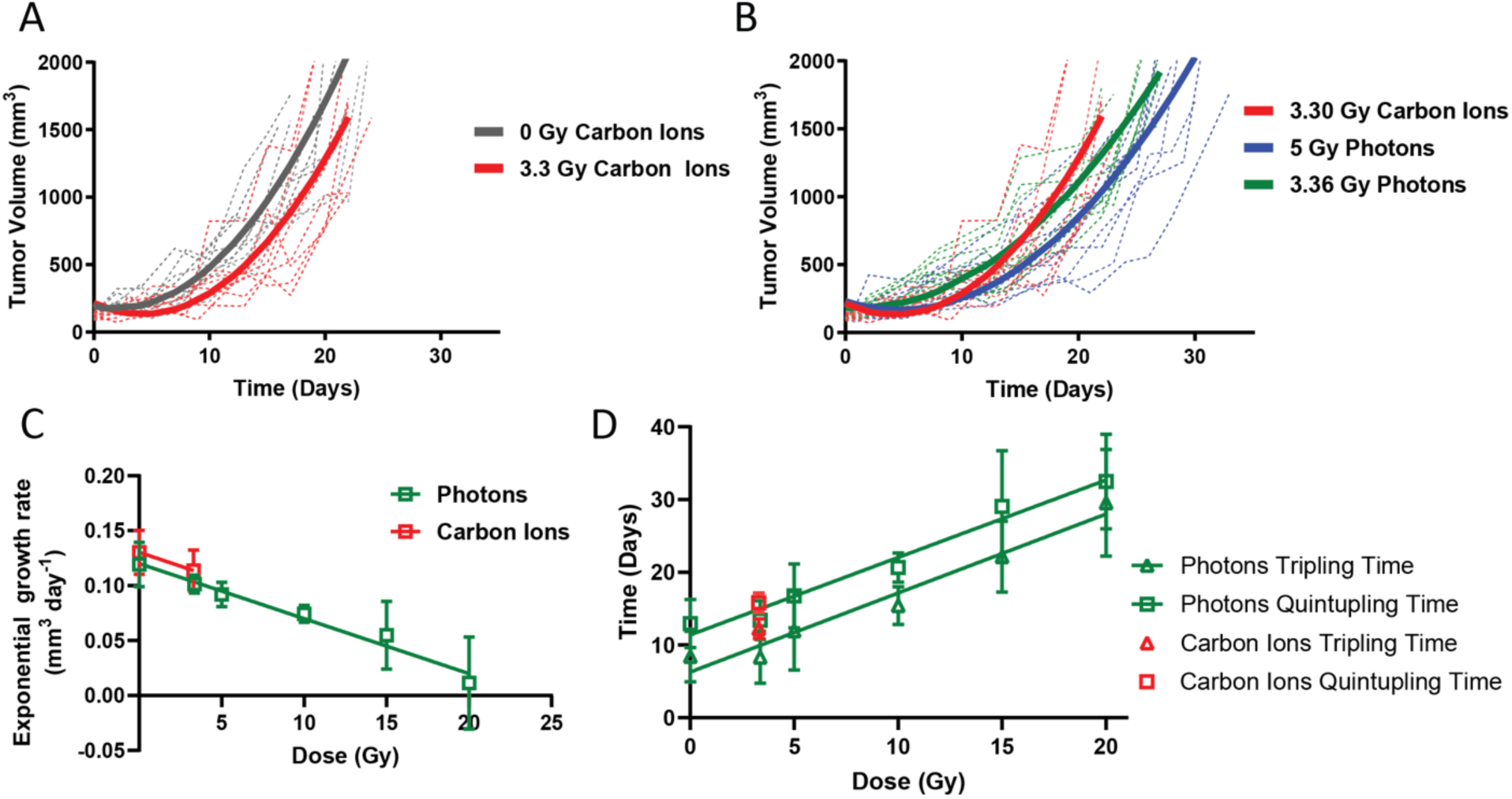
***In vivo* RBE assessment by tumor growth delay. A,** Growth curves of KPC36 tumors in control and 3.30 Gy CIRT groups; dashed lines show individual tumors, solid lines show modeled curves from repeated measures. **B,** Growth comparison among 3.30 Gy CIRT, 3.36 Gy photon RT, and 5 Gy photon RT groups. **C,** Exponential growth rates of KPC36 tumors plotted against absolute dose. **D,** Time to tumor tripling (triangles) or quintupling (squares) as a function of dose.

### Carbon ion therapy at a similar dose reveals differential immune dynamics compared to photons

To investigate the discrepancy between *in vitro* and *in vivo* RBE values, we conducted comprehensive bulk RNA sequencing analysis of KPC36 tumors following treatment with either 3.30 Gy CIRT or 3.36 Gy photon RT. Samples were collected at four-and seven-days post-irradiation to capture temporal changes in tumor transcriptomes. Using the PDAC TME classification [33], we observed significant alterations in the KPC TME four days after CIRT (Figure 5A). Compared to the combined transcriptomes of 0 Gy CIRT and 0 Gy photon controls, CIRT at day four enriched programs associated with anti-tumor immune infiltrates, including anti-tumor cytokines, cytotoxic T cells, T cells, MHC class I, co-activation molecules, and to a lesser extent, T cell trafficking and M1 macrophage signatures. Simultaneously, CIRT increased programs related to myeloid-derived suppressor cells (MDSCs) and checkpoint inhibition (Figure 5A). None of these gene signatures were elevated in the photon-treated group at day four, where instead, photon RT induced epithelial-mesenchymal transition (EMT) signatures.

**Figure 5:**
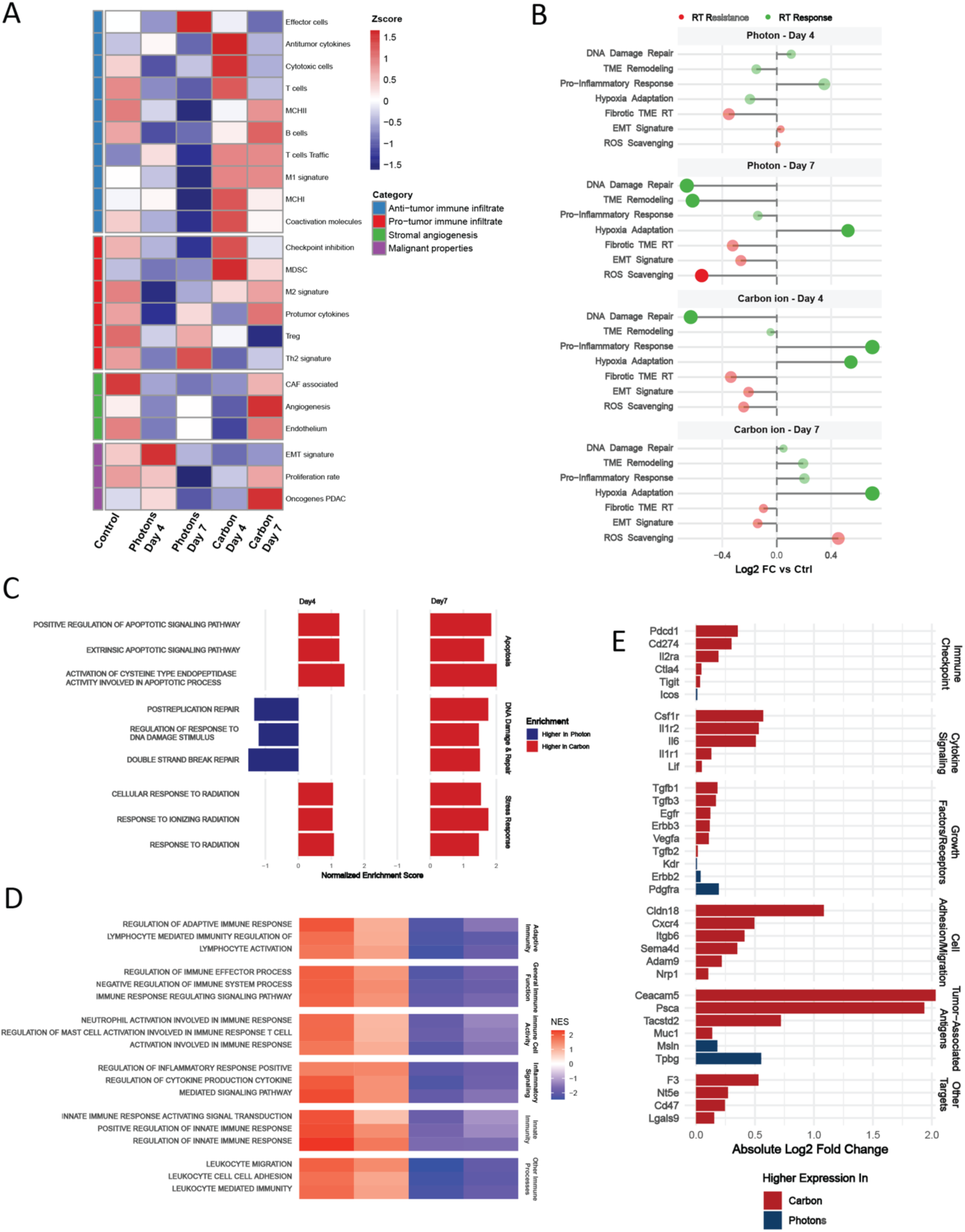
The KPC tumor immune response to carbon ion therapy differs from the response to photons. A, TME classification changes post-photon and carbon ion therapy at Days 4 and 7, D = “Depleted”, IE = “Immune enrich”, IE/F = “Immune enrich/Fibrotic”. **B,** Radiotherapy response and resistance module scores. **C,** Key radiation response pathways, shown as normalized enrichment scores (NES) & (FDR < 0.05), comparing carbon ion and photon RT at Days 4 and 7. **D,** Immune response pathways represented as NES & FDR < 0.05. **E,** Fold-change in expression of targetable **Table 1**: Surviving fraction at 2 or 3.5 Gy and RBE at surviving fraction 10% or mean inactivation dose (MID) according to either the Repairable Conditional Repairable (RCR) or Linear Quadratic (LQ) models.

In keeping with the TME classification framework [33], we observed striking alterations in the tumor immune landscape four days after CIRT (Figure 5A). CIRT-treated tumors exhibited significant enrichment of anti-tumor immune signatures, including upregulation of anti-tumor cytokines, cytotoxic T cell markers, MHC class I molecules, antigen-presenting MHC II cells, and co-activation receptors. Notably, these robust immune responses were absent in photon-treated tumors at the same timepoint, where instead, photon RT primarily induced EMT programs, effector cells, Th2 signatures, and Tregs. However, by day seven, CIRT revealed a shift toward pro-tumor pathways. This included upregulation of pro-tumor cytokines, M2 macrophage markers, PDAC oncogenes, malignant proliferation markers, and stromal components (endothelial cells, cancer-associated fibroblasts, angiogenesis factors). This temporal transition suggests that while CIRT initially stimulates potent anti-tumor immunity, while compensatory immunosuppressive mechanisms emerge by day seven while photon-treated tumors showed limited changes. Pathway analysis further delineated the differential responses between radiation modalities (Figure 5B). CIRT uniquely induced robust pro-inflammatory and hypoxia adaptation signatures at day four, with the latter persisting while inflammatory signals diminished by day seven. Notably, the transition to a pro-tumor response seven days post-CIRT was accompanied by increased expression of transcripts associated with reactive oxygen species (ROS) scavenging, suggesting that mechanisms of resistance to CIRT may involve enhanced ROS detoxification, contributing to tumor survival and progression.

Examination of radiation response pathways revealed distinct patterns between CIRT and photon irradiation (Figure. 5C). DNA double strand break repair, post-replication DNA repair, and regulation of DNA damage response pathways were more significantly enriched in photon-treated tumors four days post-irradiation. In contrast, apoptosis-related pathways were consistently more enriched following CIRT treatment compared to photons, aligning with published findings that carbon ion radiation induces higher levels of apoptosis in cancer cell lines [34–37].

We investigated immune response pathway enrichment encompassing both adaptive and innate immunity, immune cell activity, and inflammatory signaling in KPC tumors treated with either CIRT or photon RT. Our analysis revealed enhanced immune response pathway activation following CIRT compared to photon RT, most pronounced at day 4 post-treatment, which was compared to lower levels observed on day 7. In contrast, immune response pathway expression in photon-treated tumors was generally low across timepoints (Figure. 5D). These findings align with previous observations, suggesting that CIRT induces a pronounced immune response compared to photon RT [38, 39]. The disparity in immune response patterns between the two treatment modalities highlight potential benefits of CIRT for immunotherapy synergies and tumor regression.

Finally, we evaluated expression of clinically targetable genes in treated tumors (Figure 5E). In response to CIRT, we observed significant increases in the expression of cell surface glycoproteins encoded by *Ceacam5*, *Psca*, *Tacstd2*, *F3*, and *Muc1*, as well as cell adhesion proteins encoded by *Cldn18*, *Cxcr4*, *Itgb6*, and others. Additionally, CIRT upregulated cytokine signaling mediators including *Csf1r*, *Il1r2*, *Il6*, *Il1r1*, and *Lif*, as well as growth factors and their receptors, such as *Tgfb1*, *Tgfb3*, *Efgr*, *Erbb3*, *Vegfa*, and *Tgfb2*. In contrast, photon RT did not induce expression of nearly as many targetable genes. Regarding immunotherapy-related targets, CIRT uniquely increased expression of checkpoint genes like *Pdcd1*, *Cd274*, *Nt5e*, *Cd47*, *Il2ra*, *Lgals9*, *Ctla4*, and *Tigit*, while photon radiation only elevated expression of *Icos*.

These molecular insights highlight promising opportunities for CIRT-based combination treatment strategies for PDAC.

## Discussion

Our comprehensive investigation of radiotherapy response in PDAC using genetically engineered KPC mouse models provides new insights into response mechanisms and identifies potential therapeutic opportunities. The preclinical models we developed recapitulate the challenging immunosuppressive TME characteristic of human PDAC, making them valuable tools for translational research with low antigen load [40].

Based on our transcriptomic analyses, we identified several promising targets for potential CIRT combination strategies. While FAK inhibition has shown synergy with photon RT in PDAC models [41], immune checkpoint inhibitors failed to enhance response in our KPC system. However, the distinct immune activation observed after CIRT presents new opportunities. Preclinical studies in other tumor models report varied outcomes for CIRT combined with immunotherapy. For instance, αCTLA-4 with 3 Gy CIRT outperformed αPD-1 in colon-26 tumors [42], and αCTLA-4 with fractionated CIRT induced both local and distant tumor responses in EO771 mammary cancer [43]. In our KPC model, CIRT induced expression of clinically actionable targets, including PD-1, PD-L1, Claudin18, PSCA, and CEA. These findings support further investigation of checkpoint blockade and targeted agents in combination with CIRT. Such combinations may improve outcomes by leveraging the distinct immune response initiated by carbon ion therapy.

Following photon RT, KPC tumors exhibited a predominantly immunosuppressive TME characterized by increased infiltration of granulocytes, macrophages, and fibroblasts, consistent with previous reports in various cancer models [44–46]. Notably, combination immunotherapy with photon RT failed to enhance therapeutic benefit in either irradiated or unirradiated contralateral tumors, despite previous reports of potential abscopal effects in specific KPC clones [32]. The transcriptomic profile of photon-irradiated tumors aligned with the “Immune Enriched/Fibrotic” classification recently described in human PDAC [33], highlighting the translational relevance of our findings.

The discrepancy between *in vitro* and *in vivo* relative RBE of CIRT represents a central finding of our study. The RBE value for KPC36 cells based upon clonogenic survival determined at 10% survival using LQ fitting, was 3.3 while the *in vivo* RBE ranged from only 1.3-1.7 based on tumor growth delay kinetics, 3-fold vs 5-fold volume changes, respectively, the former not reflecting the full extent of tumor response. This discrepancy underscores the critical influence of the TME in modulating radiation response and highlights the limitations of relying solely on *in vitro* data for treatment planning. One likely contributor to this observed difference is our use of heterotopic tumor models. As in our study and several others [23, 38, 47], tumors were implanted in the hind limbs of mice to avoid gastrointestinal toxicity and effectively target with the carbon-ion beam, a site that may not accurately recapitulate the immune landscape or stromal interactions of orthotopic pancreatic tumors.

To further probe the biological basis of CIRT’s limited *in vivo* efficacy, we analyzed temporal transcriptomic responses. Notably, CIRT induced distinct immunologic changes compared to photon RT. At day four post-treatment, we observed upregulation of cytotoxic T cell markers, MHC class I molecules, and pro-inflammatory cytokines, suggestive of an anti-tumor immune response. These changes were absent in photon-treated tumors, which instead showed enrichment of EMT-related gene expression. By day seven, however, the immunologic profile of CIRT-treated tumors shifted toward a more immunosuppressive state, with a relative loss of anti-tumor signatures, that could be due to high proliferative rate and adaptive immune resistance.

These findings align with prior preclinical studies demonstrating CIRT-specific immune modulation. For example, CIRT has been shown to reduce infiltration of immunosuppressive myeloid cells and M2 macrophages while enhancing M1 macrophages, CD8+ T cells, and NK cells across several murine tumor models [48–50]. Despite initiating a qualitatively different immune response than photon RT, a single 3.30 Gy CIRT dose produced only modest tumor growth delay in our model. This suggests that transient immune activation alone may be insufficient to overcome the desmoplastic, immunosuppressive TME characteristic of pancreatic cancer.

In conclusion, our findings demonstrate that while the stromal architecture of PDAC attenuates the intrinsic effectiveness of CIRT, it induces unique anti-tumor immune responses that could be therapeutically exploitable. The transient nature of this immune activation suggests that appropriately timed combination strategies targeting the TME could overcome the limitations imposed by the complex PDAC microenvironment. Future studies should focus on exploiting the physical advantages of CIRT – reduced oxygen dependence, clustered DNA damage, and beam profile with low exit dose – in conjunction with immunomodulatory agents specifically targeting the dynamic immune landscape we have characterized.

## Acknowledgements

The authors would like to thank our BNL liaison, Peter Guida, as well as Delaney Felix for her assistance with animal management, and colleagues at CNAO including Angelica Facoetti and Mario Ciocca. We credit Debu Saha for assistance with IACUC protocols and the UTSW Preclinical Radiation Core Facility. Single cell RNA sequencing was performed by the Next Generation Sequencing Core, McDermott Center, UTSW. Bulk RNA sequencing was performed by the North Texas Genome Center, UT Arlington. Funds were provided by T32 funding (KLS; T32CA124334), CPRIT Recruitment of First-Time, Tenure-Track Faculty Members RR170051 (TAA), UTSW Disease Oriented Scholars Program (TAA), Carroll Shelby Family Foundation (TAA), UT Southwestern Department of Radiation Oncology Seed Grant (AJS, RH, MDS, and TAA), and David A Pistenmaa M.D. Ph.D. Distinguished Chair in Radiation Oncology (MDS).

BioRender was used to generate figures per UTSW license (Figure 1, License #*CG288WPFYF,* Figure 2, License # *CW288WRKRZ*)

## Data Availability Statement

Transcriptomic data is available at the Gene Expression Omnibus (GEO) data repository (scRNAseq GSE296677, and bulk RNAseq of treated tumors GSE296774). All other data generated in this study are available upon request.

