## Supplemental Figures for "Immune stromal components impede biological effectiveness of carbon ion therapy in a preclinical model of pancreatic ductal adenocarcinoma"

**Supplementary Materials**

**Supplementary Figure 1: KPC clonal cell lines show variable *in vivo* behavior but general therapy resistance. A,** Survival of mice bearing tumors with germline heterozygous (KPC) versus homozygous (KPPC) p53 loss. **B,** Variable *in vivo* growth of KPC clones; solid lines indicate untreated tumors, dashed lines indicate growth delay after 15 Gy photon RT. **C,** Replicating experiments by Rech *et al.* (2018) using a different cell line, KPC36 cells were injected SQ into bilateral flanks of C57BL/6 mice on Day 0 (n = 5). In photon RT-treated groups, the right sided tumor was irradiated with 20 Gy photons on Day 8. In combination immunotherapy-treated groups, αCTLA-4 and αPD-1 were given on Days 5, 8, and 11 with αCD40 given on Day 11. **D,** KPC63 cells were injected SQ into bilateral flanks of C57BL/6 mice on Day 0 (n = 5). In photon RT-treated groups, the right sided tumor was irradiated with 20 Gy photons on Day 10. In combination immunotherapy-treated groups, αCTLA-4, αPD-1, and αCD40 were given on Day 10.

**Supplementary Figure 2: Cell type origin of transcriptomic alterations in the TME after photon RT or immunotherapy in KPC tumors. A,** t-SNE distribution of tumor and tumor-draining lymph node cells in photon RT, combination immunotherapy, and control groups. **B,** Dot plot indicating gene expression patterns used to immunophenotype cells in scRNAseq data. **C,** Heatmap of transcriptional programs expressed in response and resistance modules (Supplementary Table 1) after photon RT. Color intensity indicates the relative expression level of each signature represented as z score. **D,** Heatmap of cultivated transcriptional programs expressed in response and resistance modules (Supplementary Table 1) after combination immunotherapy. Color intensity indicates the relative expression level of each signature represented as z score. **E,** Dot plot illustrating average expression of relevant targetable genes after photon RT represented in Figure 2G. Dot size represents percent of cells expressing each gene. **F,** Dot plot illustrating average expression of relevant targetable genes after combination immunotherapy represented in Figure 2G. Dot size represents percent of cells expressing each gene.

**Supplementary Figure 3: KPC36 was selected for comparison between photon RT and CIRT. A,** Comparison of cGAS (at both 5 and 45 min exposure times) and STING (at 5 min exposure) expression among KPC clones via western blot. **B,** Pilot study comparing KPC63 tumor growth from time of injection to arrival at BNL from UTSW versus tumor growth at UTSW, measured on the same dates by two separate individuals. **C,** KPC36 tumor growth in untreated control mice that were shipped to BNL alongside CIRT-treated mice and control mice that remained at UTSW alongside photon-treated mice. Dashed lines represent induvial tumors and solid lines represent average growth based on Repeated Measures Modeling. Measurements were made by the same person. **D,** KPC36 tumor growth following a range of doses of photon RT. Dashed lines represent induvial tumors and solid lines represent average growth based on Repeated Measures Modeling. **E,** Kaplan-Meier survival of KPC36 tumor-bearing mice.

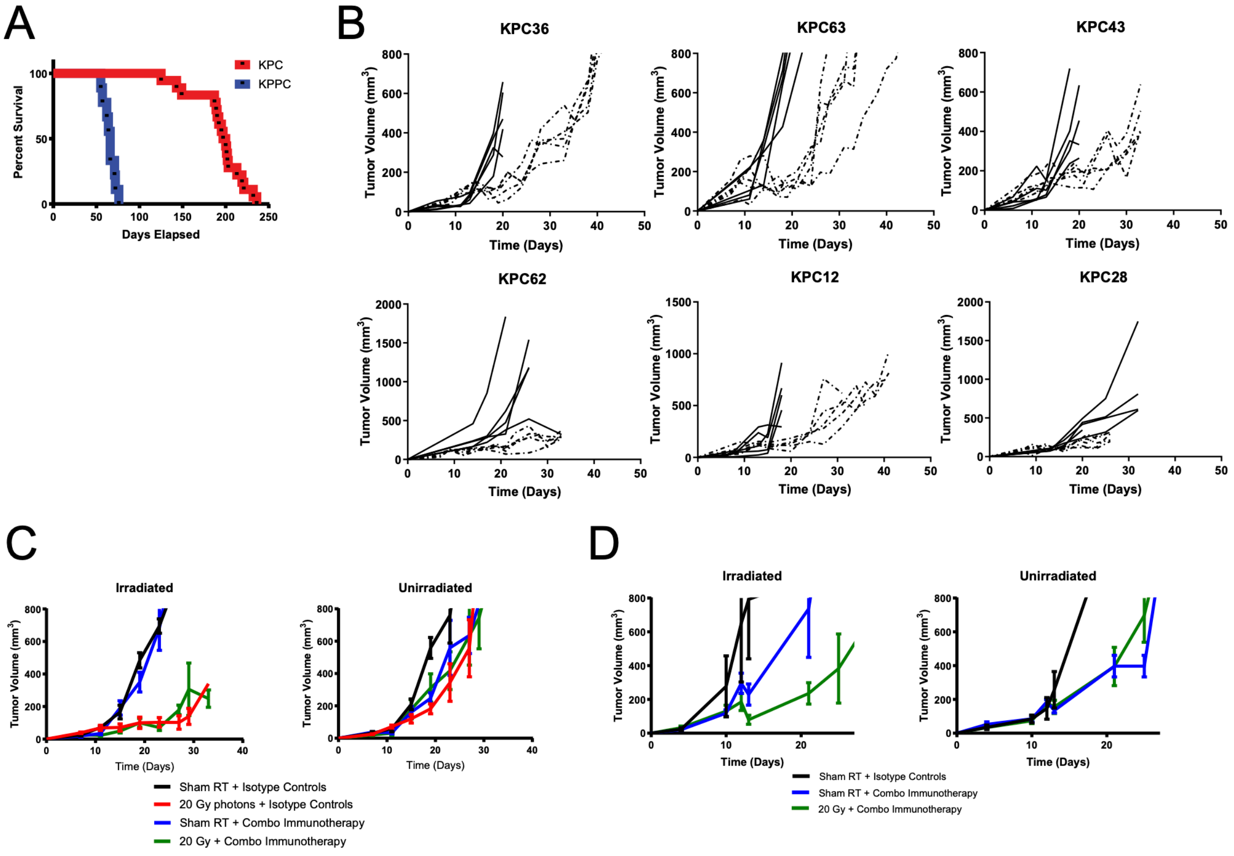


Supplementary Figure 1

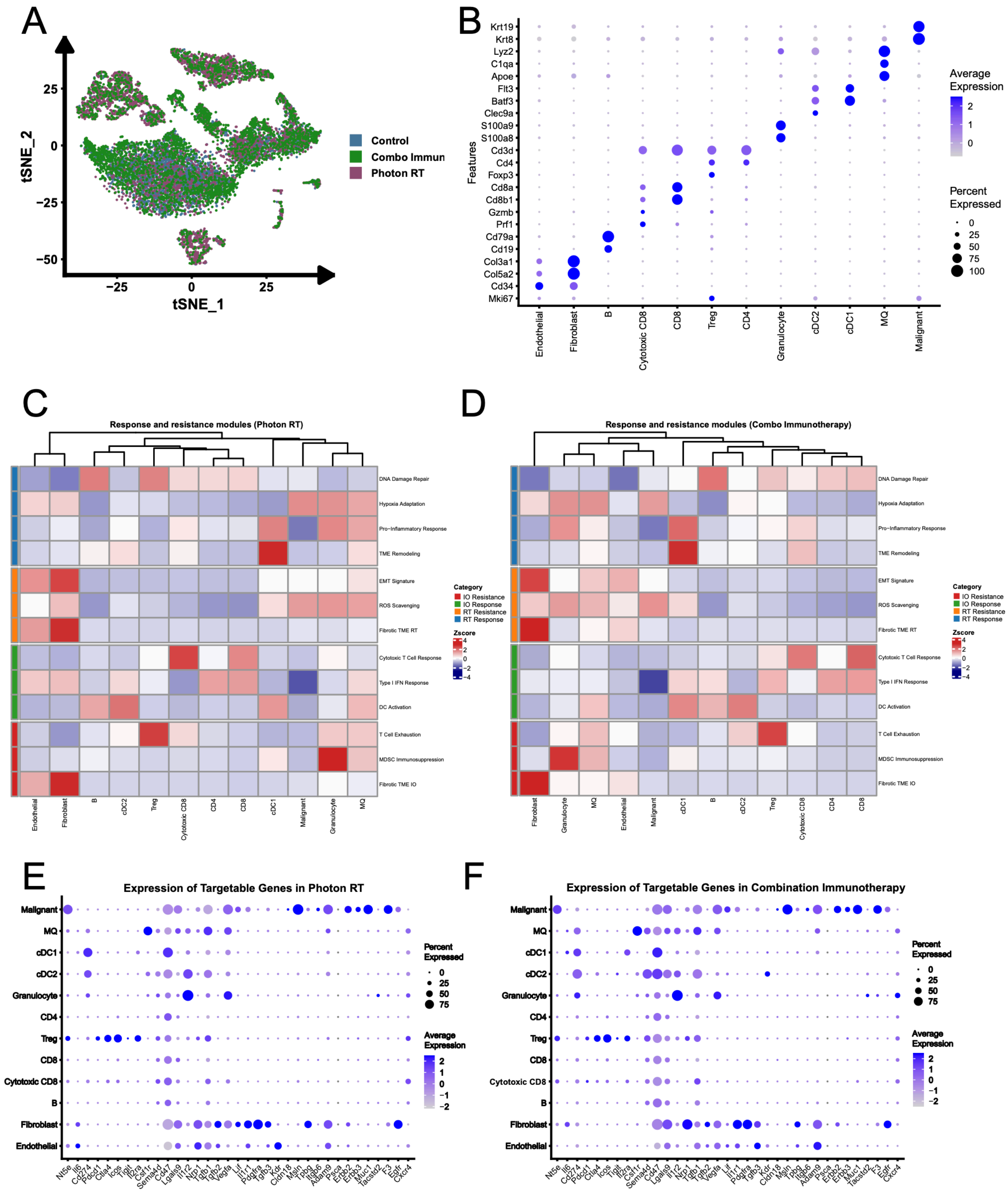


Supplementary Figure 2

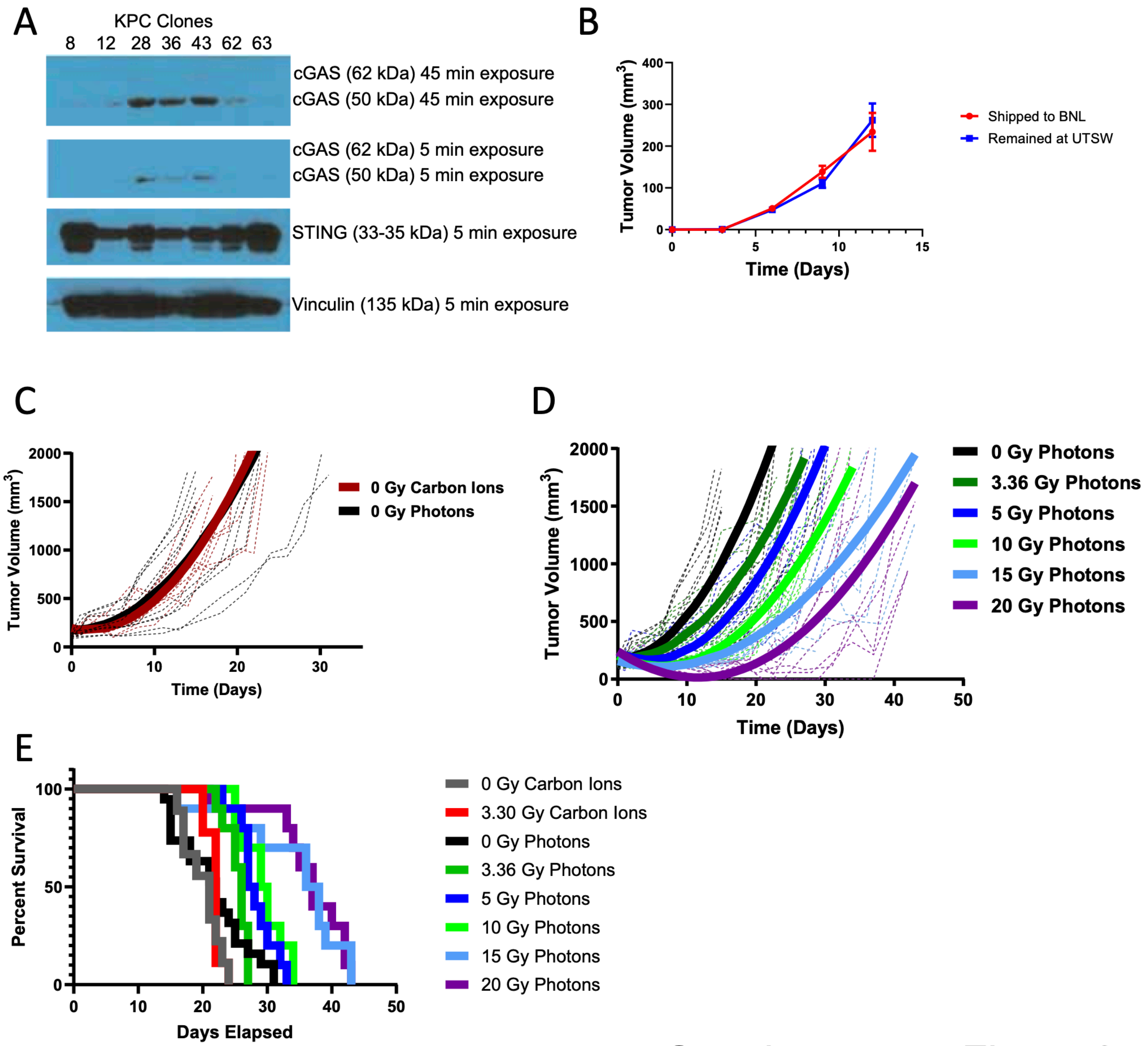


Supplementary Figure 3
